# Time-Encoding Migrates from Prefrontal Cortex to Dorsal Striatum During Learning of a Self-Timed Response Duration Task

**DOI:** 10.1101/2020.11.19.390286

**Authors:** Gabriela Chiuffa Tunes, Eliezyer Fermino de Oliveira, Estevão Uyrá Pardillos Vieira, Marcelo Salvador Caetano, André Mascioli Cravo, Marcelo Bussotti Reyes

## Abstract

Although time is a fundamental dimension of life, we do not know how the brain encodes the temporal information. Several brain areas underlie the temporal information, such as the hippocampus, prefrontal cortex, and striatum, but evidence of how they cooperate to process temporal information is scarce. Notably, the analysis of neural activity during learning are rare, mainly because timing tasks usually take a long time to train. Here we investigated how the time encoding evolves when animals learn to time a 1.5 s interval. We designed a novel training protocol where rats go from naive- to proficient-level timing performance within a single session, allowing us to investigate neuronal activity from very early learning stages. We used pharmacological experiments and machine-learning algorithms to evaluate the level of time encoding in the medial prefrontal cortex and the dorsal striatum. Our results show a double dissociation between the roles of the medial prefrontal cortex and the dorsal striatum during temporal learning, where the former commits to early learning stages while the latter become more engaged as animals become more proficient in the task.

## Introduction

Even though keeping track of time is essential for survival, our understanding of how animals encode such information in terms of neuronal activity is still modest^1, 2^. The literature contains evidence of physiological activity accounting the temporal information, as for example ramping-neurons^3–5^ and time-cells^6, 7^, neuronal oscillations^8^, sequential firing of neurons^9, 10^. Most of these encoding patterns are particular cases of a general type of encoding called population clocks, which states that any reliable dynamical evolution of neural activity works as a potential clock and might serve as a timing mechanism^11^. However, establishing causal links between the time encoding and timing itself has been elusive^9, 12^.

Reports of the spiking activity of neurons involved in keeping track of time implicate regions like medial prefrontal cortex^5, 8, 13–15^, motor cortex^10, 16, 17^, striatum^2, 5, 8, 10, 18–20^, hippocampus^6, 7^, thalamus^21^, substantia nigra pars compacta^22^, among others. Active manipulations of the physiological activity have also helped assess the involvement of these areas in timing, particularly in the medial prefrontal cortex^5, 22–24^ and striatum^20^. However, because timing tasks usually require many training sessions, the neurophysiological process underlying their acquisition has been less studied. The vast majority of the electrophysiological recordings in timing tasks come from very well-trained animals (but see^9^). Critically, observing how activity in different regions evolves as a function of temporal learning can improve the understanding of brain mechanisms underlying this ability.

We investigated how spiking activity in the medial prefrontal cortex (mPFC) and striatum (STR) encodes a time interval during learning. We trained rats to sustain responses for at least 1.5 s and investigated how well the two brain areas decode the time interval during acquisition. If a brain area is involved in the encoding of that interval, a trial-dependent and more structured spiking activity must emerge due to learning. As a consequence, the performance of algorithms that evaluate the decoding should increase. Contrarily to evidence of the concomitant involvement of mPFC^23, 24^ and STR^18^ in timing tasks, our results show very different roles of these two regions during learning. As training progresses, the decoding performance based on electrophysiological recordings decreases in the mPFC while it increases in the STR. Such results were confirmed and further investigated with pharmacological experiments. Our studies provide both a useful method to investigate electrophysiological correlates of temporal learning and also advance our understanding of the role of mPFC and STR in interval timing.

## Results

### Rats learn to time in a single session

We conceived a novel experimental design in which rats improve their timing in a single session^25^, allowing us to track the activity of individual neurons during learning (Fig. 1). Animals had to remain in a nose poke for at least 1.5 s to receive access to the sucrose solution, limited to three licks, after which the access gate closes (Fig. 1A). Shorter responses produced no consequences. Critically, we recorded activity before, during, and after animals had learned the critical interval.

**Figure 1.**
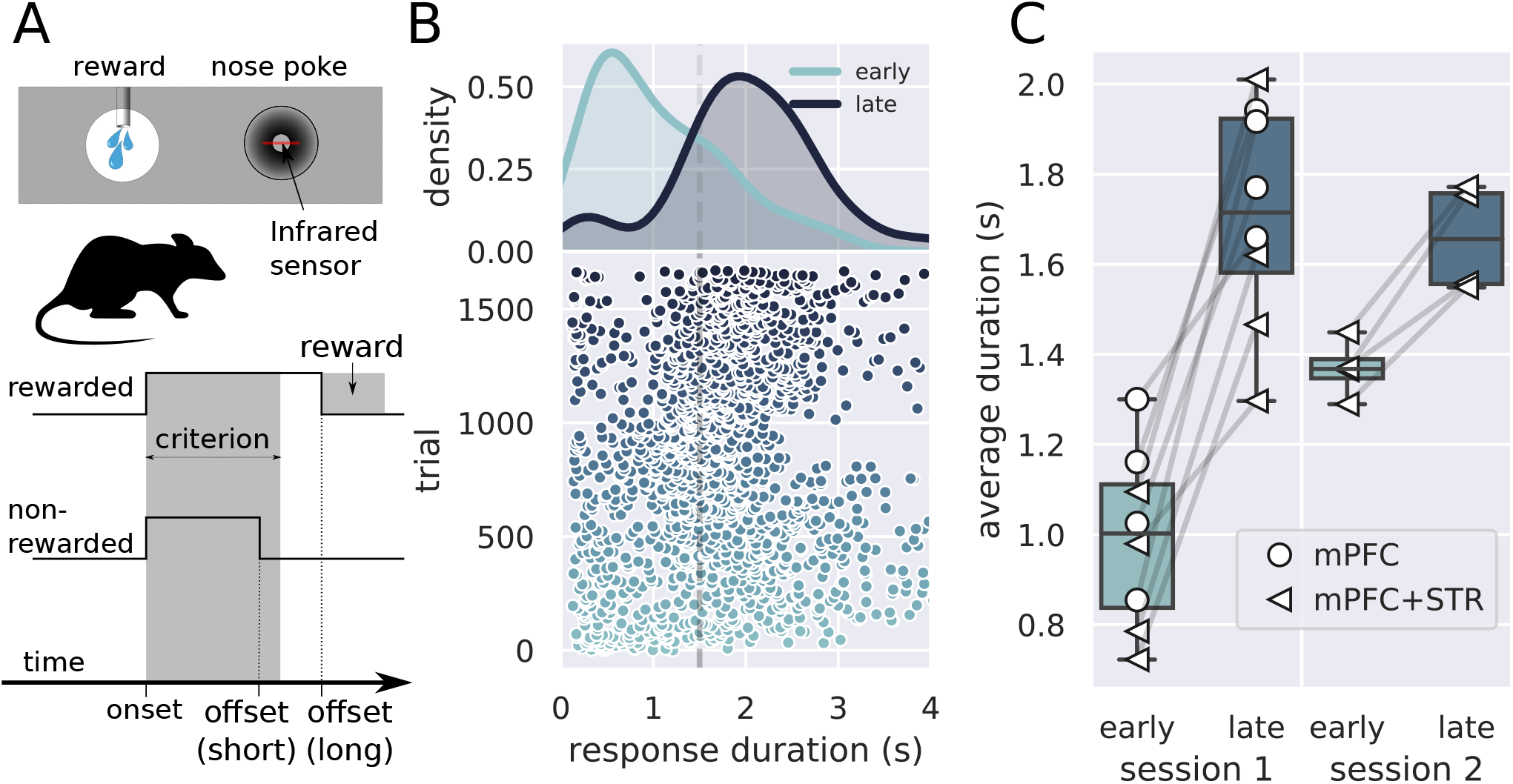
Description of the task and behavioral results. A) Experimental design used in the experiment. The trial started when the rat inserted its snout in the nose poke installed next to the reward port. The step-lines represent the nose poke in and out. Depending on how long the rat stays in the nosepoke — the response duration —, it may receive access to a sucrose solution. Responses lasting more than 1.5 s granted access to three licks of sucrose solution, while shorter responses produced no consequences. B) Responses from a single rat in the first session of training in the task. (Lower graph) Each dot represents a response duration (x-axis) in a specific trial (y-axis). The dashed line represents the criterion (1.5 s) for receiving the reward. (Upper graph) Probability density of responses as a function of the duration for early and late trials, showing that the distribution shifted to the right. C) Mean response duration for individual rats early and late in sessions 1 and 2. All animals significantly increase their responses in the very first session of training. For one group of animals (group mPFC+STR), we recorded a second day of training. At the beginning of this second session (early trials) animals’ response durations were shorter than the late trials of session 1, but much longer than the early trials, showing that the training from session one improved the behavior early in session 2. Throughout the second session, rats achieve the same performance as the late trials of session 1, suggesting that the training in the first session took the rats close to the optimal performance.

We implanted two sets of animals, one with electrode arrays in the mPFC (group mPFC, N=4 animals) in long sessions (*>* 4 hours) and another with arrays both in the mPFC and STR (group mPFC+STR, N=4 animals) in shorter sessions. Animals produced between 801 and 1671 trials (878 trials on average) in long sessions (group mPFC) and, in shorter sessions (group mPFC+STR), between 436 and 936 (606 on average) on day 1 and between 381 and 699 (535 on average) trials on day 2.

All eight animals showed significant learning in the first session, producing longer lever presses *T* and yielding higher reward rates late at the session (*t*(7) = 7.9, *p* = 10^−4^, cohen-d= 3.2, 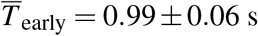, 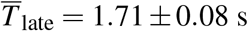; Fig. 1B and C).

Animals who trained for two days (group mPFC+STR, N=4) also produced longer responses late in the first session as compared to the beginning (Fig. 1C, group mPFC+STR). In this second session, the rats’ responses were initially shorter than those at the end of session one, increasing throughout the session. Hence, animals retained information from session 1 to 2. Furthermore, rats performed similarly at the end of both sessions, suggesting that animals learned the task almost to their best on the first day of training. A repeated measure ANOVA showed an effect of moment (*F*_[1,3]_ = 61.0, *p* = 0.004, 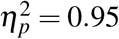), no effect for day (*F*_[1,3]_ = 2.37, *p* = 0.22, 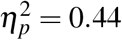) and an interaction moment versus day (*F*[1,3] = 26.0, *p* = 0.014, 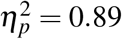). A posthoc analysis revealed a significant difference between early and late responses in the first (*t*(3) = 7.02, *p* = 0.006), but not in the second session (*t*(3) = 2.90, *p* = 0.06).

### Time representation decreases with learning in the medial prefrontal cortex, but not in the striatum, in the first day of training

We investigated the progression of individual neurons’ activity in the medial prefrontal cortex (mPFC) and the dorsal striatum (STR) during learning (Fig. 2). We examined how the neuronal activity in both regions modulated during trials and how such modulation evolved with learning (Fig. 2C–F for mPFC and G–J for STR).

**Figure 2.**
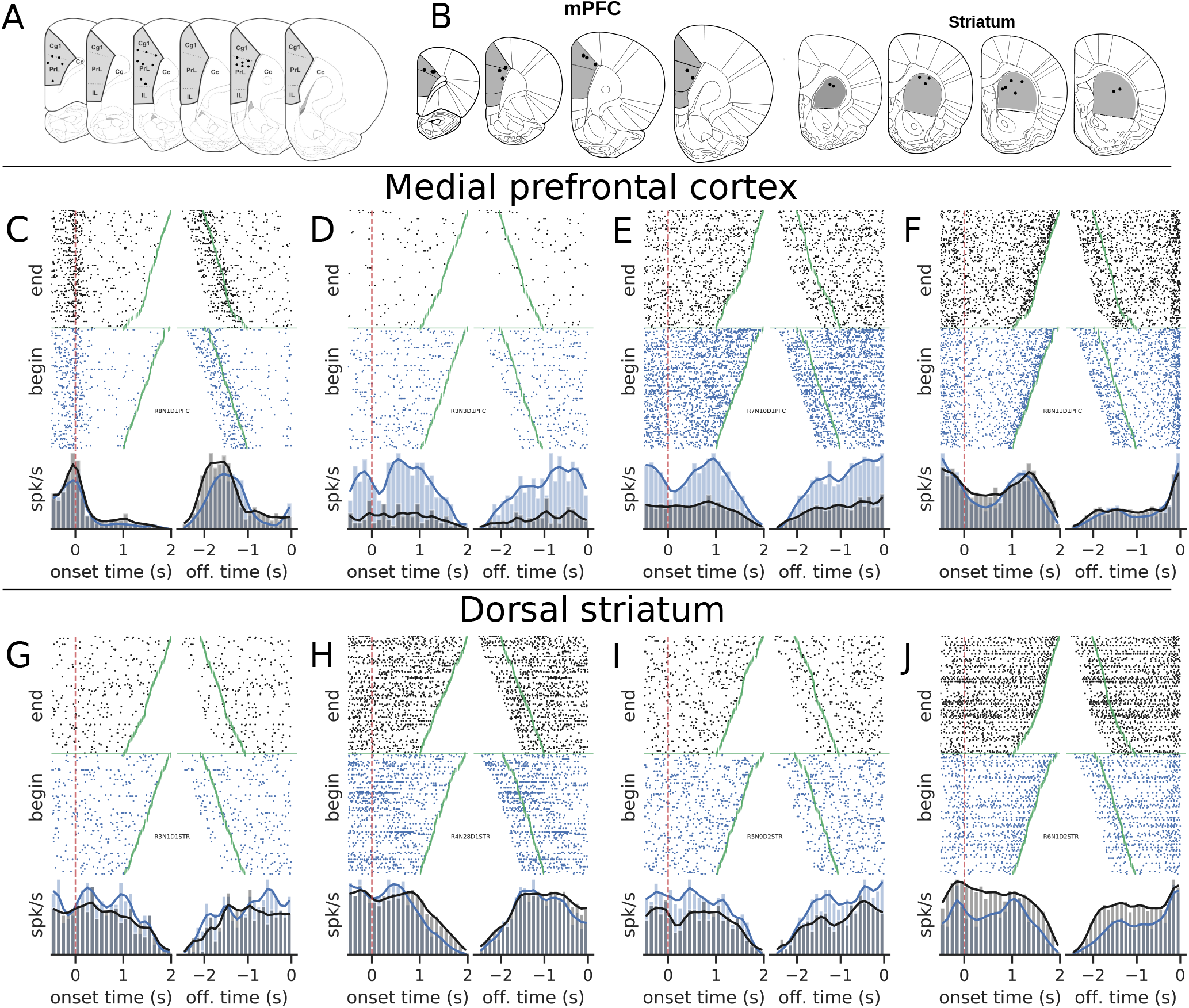
Examples of neural activity modulated during trials. A) Recording sites in the medial prefrontal cortex (group mPFC). B) Recording sites for group mPFC+STR. C–J) Upper graphs show the raster plots from early (blue) and late (black) trials for mPFC (C–F) and STR (G–J). Bottom graphs show the corresponding firing rates, color-coded. C) A neuron whose activity is tuned to the trial onset. D to F) Examples of neurons whose activity within the trial became less sensitive (flatter) late compared with early trials of the session.

Neurons exhibited diverse activity patterns, with a small number of units modulated by the onset or offset of the nose poke and others that became less responsive to trials during learning. Notably, in the mPFC, some neurons’ activity climbed up or down during the initial trials but became flat late in training. Overall, there was no evidence that the climbing activity increased with training, neither in the mPFC (*t*(3) = 0.54, *p* = 0.60, 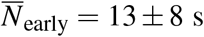, 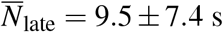) or STR (*t*(3) = −0.081, *p* = 0.94, 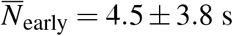, 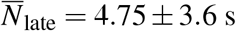). These results were at odds with our working hypotheses that the mPFC and the STR were highly involved in keeping track of time and that, once learning took place, an encoding scheme should emerge. However, the evidence for mPFC data pointed to a very distinct scenario: a vanishing neural code.

Given the heterogeneity of individual neurons’ responses and based on recent proposals of population clocks, we explored whether the pattern of activity in the mPFC and STR could work as a stable representation of time (Fig. 3. We used spikes from all recorded neurons during reinforced trials, i.e., trials longer than the criterion (1.5 s) and shorter than 3.5 s. We then truncated all trials at 1.5 s and calculated the firing rate in bins of 100 ms. This procedure results in a matrix of firing rates, representing the discrete bins, for all neurons, in all trials. We used these data to train a machine-learning model (linear discriminant analysis, LDA) to decode the time bin using firing rates as inputs and measured the classifier’s performance as the correlation coefficient between the predicted bin and the actual time-bin. We calculated Pearson’s R using 100 different training and testing sets, generating a distribution of R values compared between groups and conditions.

**Figure 3.**
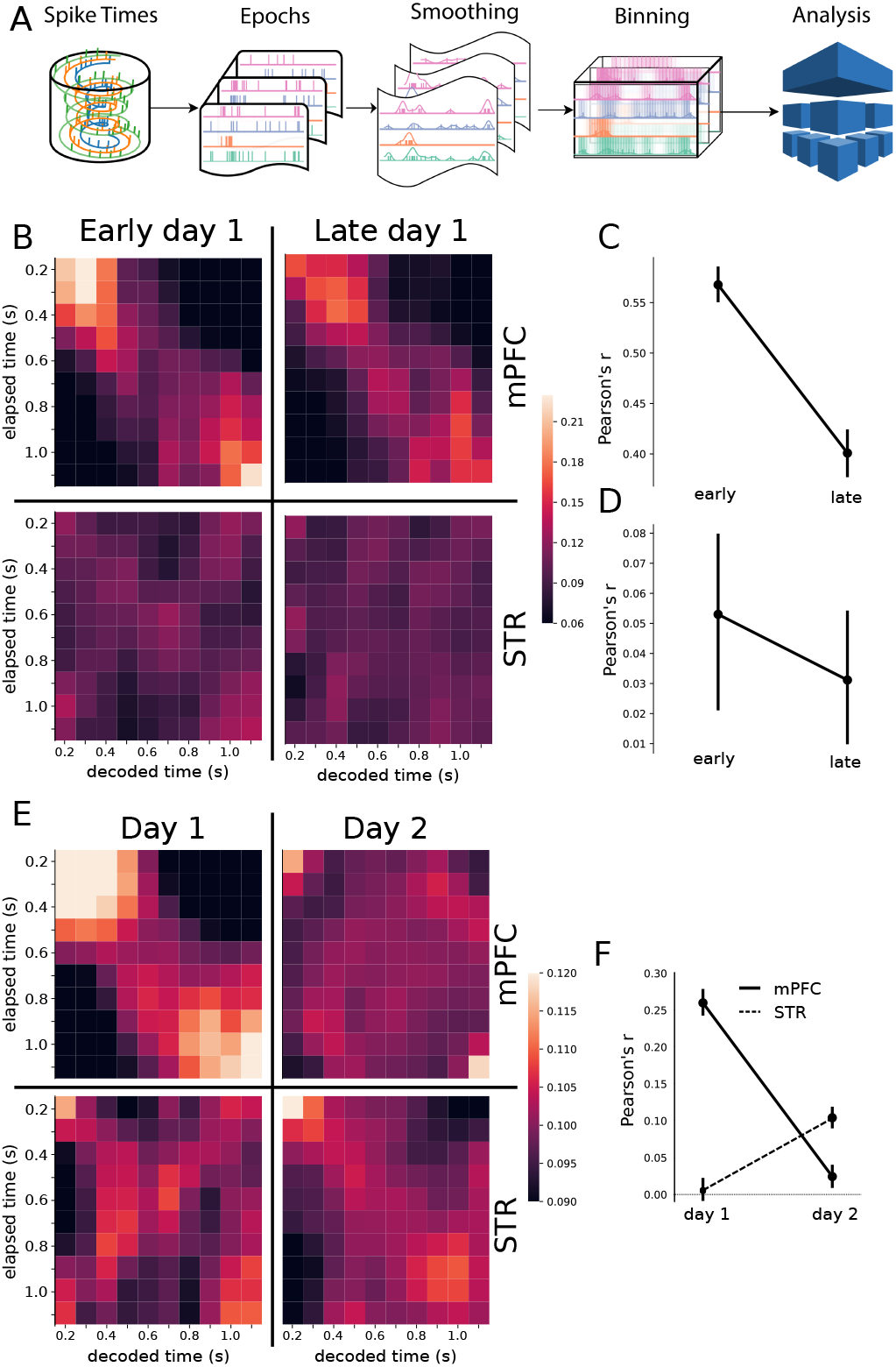
Classification analysis. A) Pipeline of the analysis at the population level. Spike trains are epoched according to the trials, from 0.5 s before nose poke onset through the offset. The epoched trials are shown in 2C–J. The trials are smoothed with a Gaussian kernel (*σ* =100 ms) and then binned every 100 ms. The resulting spike rates were analyzed with the linear discriminant analysis. B and E) Results of the classification decoder analysis within session 1 (B) and between sessions 1 and 2 (E). The confusion matrix (in the form of a heatmap) shows the probability of decoding a particular actual time bin (x-axis) as one of the possible decoding bins (y-axis). A diagonal pattern of bright pixels indicates a better performance of the classifier. B) Results of the classifier in the mPFC (top row), early and late in training showing that the performance decreased with learning. C) Overall classifier performance for mPFC data, measured as a correlation coefficient (Pearson’s r) between the actual and the predicted bin. The graphs at the bottom row in (B) show the same analysis for STR within session 1, and (D) shows the overall classification performance of this region. The results show that the performance decreases in the mPFC while there is no evidence of changes in the STR. E–F) Same analysis as in B–D but comparing data from sessions 1 and 2, for animals with electrodes implanted in both regions (group mPFC+STR). E) Decoding performance from PFC (top row) and STR (bottom row) during day 1 (left column) and day 2 (right column). During day one, PFC outperformed the STR in decoding time information. On day 2 the opposite pattern emerged. F) Pearson’s R score of the decoder performance summarizing results shown in E.

The classifier reliably decoded the temporal information from mPFC neurons (Fig. 3B-C), both early and late in session 1 (Pearson’s *R*_early_ = 0.475±0.007, t(99)=67, p=10^−84^, *R*_late_ = 0.437±0.005, t(99)=74, p=10^−88^), suggesting that the this region encoded the temporal information from very early stages of training.

The classifier revealed a distinct scenario for the STR in the first session. Even though the decoder was able to retrieve information from striatal neurons, i.e. the R-values were significantly different from zero (Pearson’s *R*_early_ = 0.066±0.012, t(99)=5.12, p=8×10^−7^, *R*_late_ = 0.046±0.013, t(99)=, p=0.0006), the decoding power was much smaller than in the mPFC, either at the beginning or at the end of the first session (Fig. 3B, lower row). These results show that the prefrontal cortex exceeded the striatum in time decoding in session 1.

We quantified how different the two regions encoded time using a two-way ANOVA comparing R values’ distributions as a function of the moment (early versus late) and region (mPFC versus STR). This analysis showed a significant effects for region (*F*_[1,99]_ = 1437, p=8×10−134, 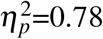), moment (*F*_[1,99]_ = 59, p=10−13, 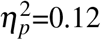) and an interaction ((*F*_[1,99]_ = 34, p=10−8, 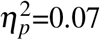, two-way ANOVA). A posthoc analysis showed significant differences between mPFC and STR, both early and late in the first session (all p-values *<* 10^−3^), confirming that the encoding level of the mPFC outperformed the STR in absolute values. Such results, however, should be taken with caution because the number of neurons recorded in the mPFC was greater than in the STR, a factor that affects the encoding performance.

Beyond the classifier’s absolute predicting power, affected by the number of neurons recorded, we were interested in how the performance changed in each region as the animals learned in the first day of training — going from a naive animal to almost a proficient level. Better performance in late trials would reveal that the region developed a time-related neural code.

The interaction between moment and region revealed that the mPFC and STR evolved differently in the session. The performance of the classifier in the mPFC decreased with training (Holm’s post hoc, p=10^−5^, Fig. 3C–D), reinforcing the previous evidence that neurons in the mPFC disengaged from timing as the training progressed. However, no difference between STR early versus late (Holm’s posthoc, p=0.26) suggests no difference between the decoding performance at the beginning and the end of the session.

We further analyzed the decoder performance in the two brain regions during two consecutive training days (Fig. 3E-G). For these experiments, we only used rats from group mPFC+STR, for which all rats had simultaneous recordings in both brain regions.

Confirming the tendency seen in the first session, the performance decreased in the mPFC from the first to the second session. On the other hand, the decoding performance in the STR neurons — which was almost negligible in the first session — significantly increased by the second session. A comparison between the classifier performance revealed a strong effect for region (*F*_[1,99]_ = 98, p=9×10−21, 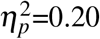, day (*F*_[1,99]_ = 110, p=7×10−23, 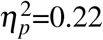) and an interaction region-day (*F*_[1,99]_ = 374, p=4×10−59, 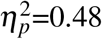). A posthoc analysis showed that during the first session, the mPFC outperformed the STR (p=5×10^−43^), but the scenario inverts during session 2, where STR neurons take over, exceeding mPFC in decoding time (p=7×10^−14^). Such an effect reveals the complementary roles of the mPFC and STR in learning.

### Medial prefrontal cortex is crucial for the acquisition but not the expression of timed responses

Given the unexpected finding of how training modulated activity in the mPFC, we further investigated its role in learning, looking for causal evidence supporting the results obtained with electrophysiological recordings. We designed a 5-day training protocol to assess the temporal performance in the first days of training with the mPFC under muscimol inactivation. We used the behavioral task shown in Fig. 1A, except that the rats responded to a lever instead of a nosepoke. In the three first sessions, the experimental group (Muscimol) received muscimol (100 ng/0.5 *μ*l) infusion in the mPFC, while the control animals (group Saline) received saline (0.5 *μ*l). On the fourth day, both groups received saline, and in the fifth session, both received muscimol.

The mPFC inactivation severely impaired learning, but had no effect on the performance of trained animals (Fig. 4). Although both Saline and Muscimol rats produced short responses early in training, only rats in the Saline group made longer responses as the session progressed (Fig. 4B). In the first session, a mixed ANOVA showed an effect for group (*p* = 0.048, *F*_[1,17]_ = 4.5, 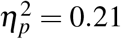), moment (*p* = 0.0006, *F*_[1,17]_ = 17.5, 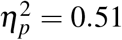), and an interaction between group and moment (*p* = 0.0049, *F*_[1,17]_ = 10.5, 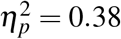). A Holm’s posthoc correction showed that the groups differed late (*p* = 0.023) but not early in the session (*p* = 0.91). Also, the group Muscimol displayed no significant difference in the response duration comparing early versus late trials (*t*(7) = 0.36, *p* = 0.73, paired t-test), with an anecdotal Bayes factor favoring the null hypothesis (*BF*_10_ = 0.36). Therefore, the group Saline learned in the first session, while the group Muscimol displayed no sign of change in behavior.

**Figure 4.**
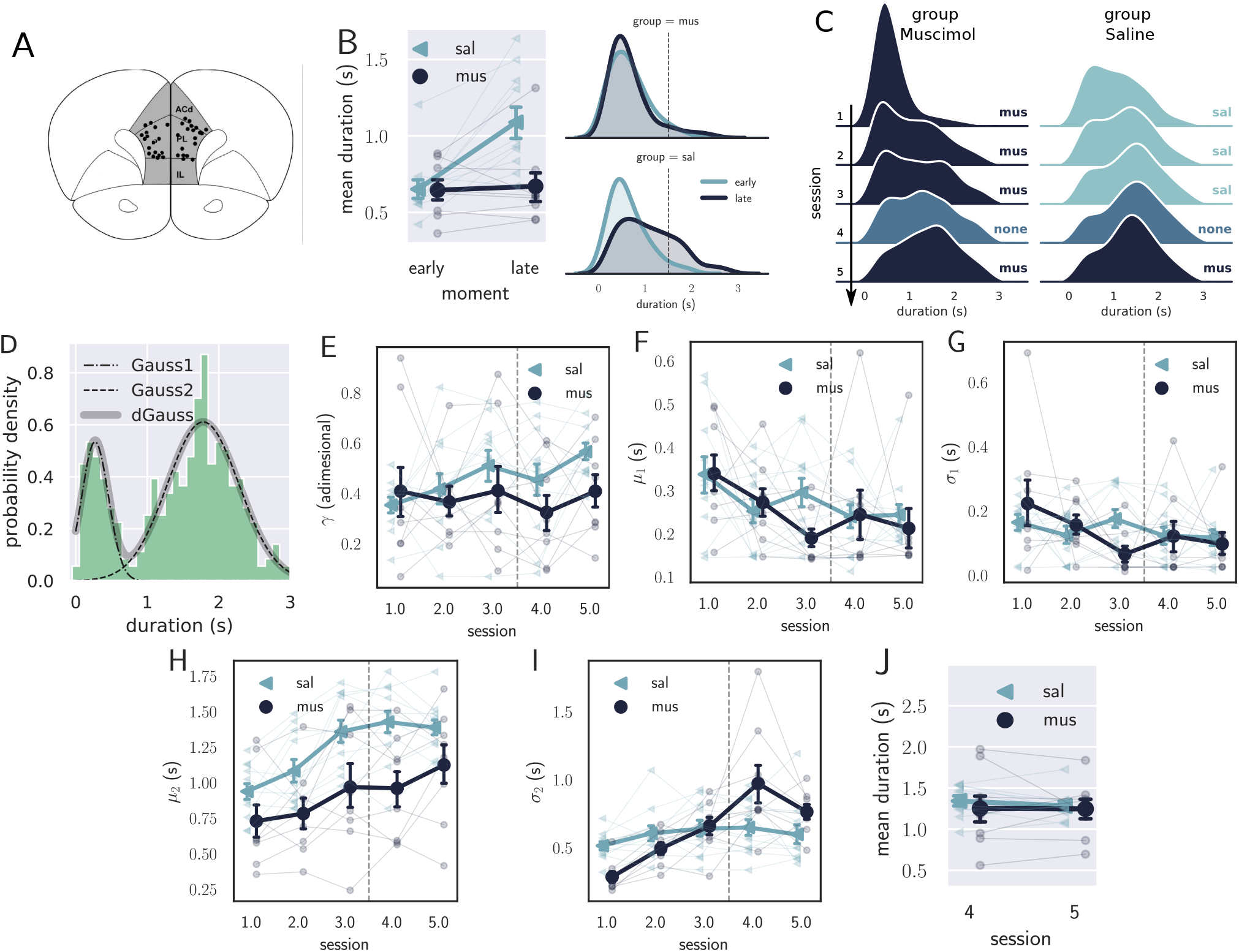
Results from the pharmacological inactivation of the mPFC. A) Histology showing the microinjection sites in the mPFC. B) Left: Average response duration early and late in the first session for group Saline (sal) and Muscimol (mus). Right: Same data as in left, but combining the responses from all rats for each group. The group Saline (sal) increased the response duration during the first session (day 1), while the group muscimol (mus) produced no detectable change in behavior. C) Probability density of late responses combined from all rats for the five consecutive sessions. The drug injected in each session is at the right of each graph and color-coded. The group Saline evolves as expected, producing longer responses from the first session, producing bimodal distribution, whose second peak increases with respect to the first, remaining stable in sessions 3 and 4. The group Muscimol has no sign of learning in the first session but progressed after the second session at a slower rate compared to group Saline. By session 3, the peak of premature responses is still more prominent than the long (*>* 1 s) responses. D) Histogram of response durations from one rat showing the bimodal distribution and the separate fitted distributions representing the premature responses (Gauss1), the temporally controlled responses (Gauss2) and the double gaussian fit (dGauss). E-I) Evolution of each double-Gaussian parameter as a function of the sessions, *γ* (E), *μ*_1_ (F), *σ*_1_ (G), *μ*_2_ (H), and *σ*_2_ (I). The vertical dashed line divides the experiment into two phases: the first (sessions 1 to 3), when groups received different treatments, and second (sessions 4 and 5) when both groups received the same treatment. J) Comparison between sessions 4 (no drug) and 5 (muscimol) for the group Saline (sal) and Muscimol (mus), suggesting that the muscimol injection produced no effect on the experienced animals.

The effect of mPFC inactivation persisted over the first three sessions impairing learning (Fig. 4C). Even though both groups learned over the three sessions, shown by an effect of session (*F*_[2,34]_ = 20.8, *p* = 10^−6^, *η*^2^ = 0.55, mixed ANOVA), rats from group Saline produced longer responses compared with the group Muscimol (effect of group, *F*_[1,17]_ = 8.27, *p* = 0.01, *η*^2^ = 0.32), with no significant interaction (*F*_[2,34]_ = 0.29, *p* = 0.75, *η*^2^ = 0.017).

To further investigate the underlying changes induced by the mPFC inactivation, we fitted the distributions of response durations with a double Gaussian function (Fig. 4D). The distributions became bimodal during learning, and it has been shown25 that such a phenomenon happens not only at the group level — what could suggest an artifact of group-averaging —, but also at the individual level. Even well-trained animals display bimodal distributions, suggesting that rats alternate between responses of two classes: premature (short) and time-controlled (long, Fig. 4 D). We investigated how the two classes of responses evolve using the five parameters that the double-Gaussian yields: *γ* ∈ [0, 1], *μ*_1_, *σ*_1_, *μ*_2_, and *σ*_2_ (*μ*_1_ ≤ *μ*_2_). The parameters *μ* and *σ* are the mean and the standard deviation of the Gaussian distributions, and *γ* represents the ratio between their amplitude. We adjusted a double-Gaussian curve for each animal and session and plotted the extracted parameters as a function of the session.

The double-Gaussian analysis confirmed that learning was impaired in the group Muscimol and revealed that the effect was mainly in the temporally-controlled responses (Fig. 4E-I). Over sessions, although the temporal-controlled responses (mean of the second Gaussian, *μ*_2_) became longer for both groups, this increase was larger for the saline group (main effect of group, *F*_[1,17]_ = 7.12, *p* = 0.016, *η*^2^ = 0.30; main effect of session:*F*_[2,34]_ = 11.20, *p* = 0.020, *η*^2^ = 0.40; interaction: *F*_[2,34]_ = 0.72, *p* = 0.49, *η*^2^ = 0.04). The standard deviation the temporally controlled responses (*σ*2) also evolved differently in the saline group (main effect of group, *F*_[1,17]_ = 3.21, *p* = 0.09, *η*^2^ = 0.16; main effect of session:*F*_[2,34]_ = 17.05, *p <* 0.001, *η*^2^ = 0.50; interaction: *F*_[2,34]_ = 4.82, *p* = 0.04, *η*^2^ = 0.22).

The premature responses (associated with the first Gaussian) became shorter for both groups, i.e. *μ*1 decreased (main effect of group, *F*_[1,17]_ = 0.70, *p* = 0.41, *η*^2^ = 0.04; main effect of session:*F*_[2,34]_ = 4.16, *p* = 0.024, *η*^2^ = 0.19; interaction: *F*_[2,34]_ = 2.05, *p* = 0.14, *η*^2^ = 0.11). Neither *σ*_1_ or *γ* were significantly modulated across the first three sessions (*σ*_1_: main effect of group, *F*_[1,17]_ = 0.05, *p* = 0.81, *η*^2^ = 0.003; main effect of session: *F*_[2,34]_ = 1.45, *p* = 0.27, *η*^2^ = 0.78, interaction: *F*_[2,34]_ = 2.76, *p* = 0.08, *η*^2^ = 0.14; *γ*: main effect of group, *F*_[1,17]_ = 0.20, *p* = 0.66, *η*^2^ = 0.01; main effect of session:*F*_[2,34]_ = 1.33, *p* = 0.28, *η*^2^ = 0.07, interaction: *F*_[2,34]_ = 0.86, *p* = 0.42, *η*^2^ = 0.04). Overall, these differences point to decreased temporal control when the mPFC is inactivated.

The last phase of the experiment (sessions 4 and 5) aimed at investigating the effect of the mPFC inactivation on experienced animals. By session 4 — when rats received no injections —, the rats from both groups were well trained, mostly producing temporally-controlled responses. We compared the results from sessions 4 (no injection) and 5 (muscimol) to check how the inactivation of the mPFC affected the response distributions after learning.

The results from the last phase showed that inactivating the mPFC after training produced no detectable effect on the response distributions (Fig. 4J). A mixed ANOVA yielded no significant effect for session (*F*_[1,17]_ = 0.09, *p* = 0.77, *η*^2^ = 0.005), group (*F*_[1,17]_ = 0.55, *p* = 0.47, *η*^2^ = 0.03) or interaction between the factors (*F*_[1,17]_ = 0.033, *p* = 0.86, *η*^2^ = 0.002). Such a result indicates that the mPFC is only required at the beginning of the task — during the learning phase—, becoming unnecessary as the animals get proficient. This finding is consistent with our previous observation regarding the disengagement of the mPFC observed in the electrophysiological recordings within the first session.

### Striatum is necessary for temporally-controlled responses

Results from the decoding analysis also suggested a higher participation of the STR in the temporal task in the second training session. To confirm this finding, we inactivated the STR in experienced animals, after four sessions of training. STR inactivation strongly impaired temporal performance, leading to a lower mean response duration (*t*(3) = 5.6, *p* = 0.011, cohen-d=0.92, two-tailed t-test, Fig. 5B–C). We again fitted double Gaussians and compared their parameters across sessions four and five (before and during STR inactivation, respectively, Fig. 5D–H). We observed no significant effect for gamma (*t*(3) = 2.3, *p*_*γ*_ = 0.11), *μ*_1_ (*t*(3) = −0.38, 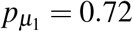), or *σ*_1_ (*t*(3) = 2.6, 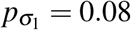), but significant decrease in *μ*_2_ (*t*(3) = 3.36, 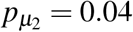), and an increase in *σ*_2_ (*t*(3) = −5.66, 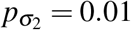).

**Figure 5.**
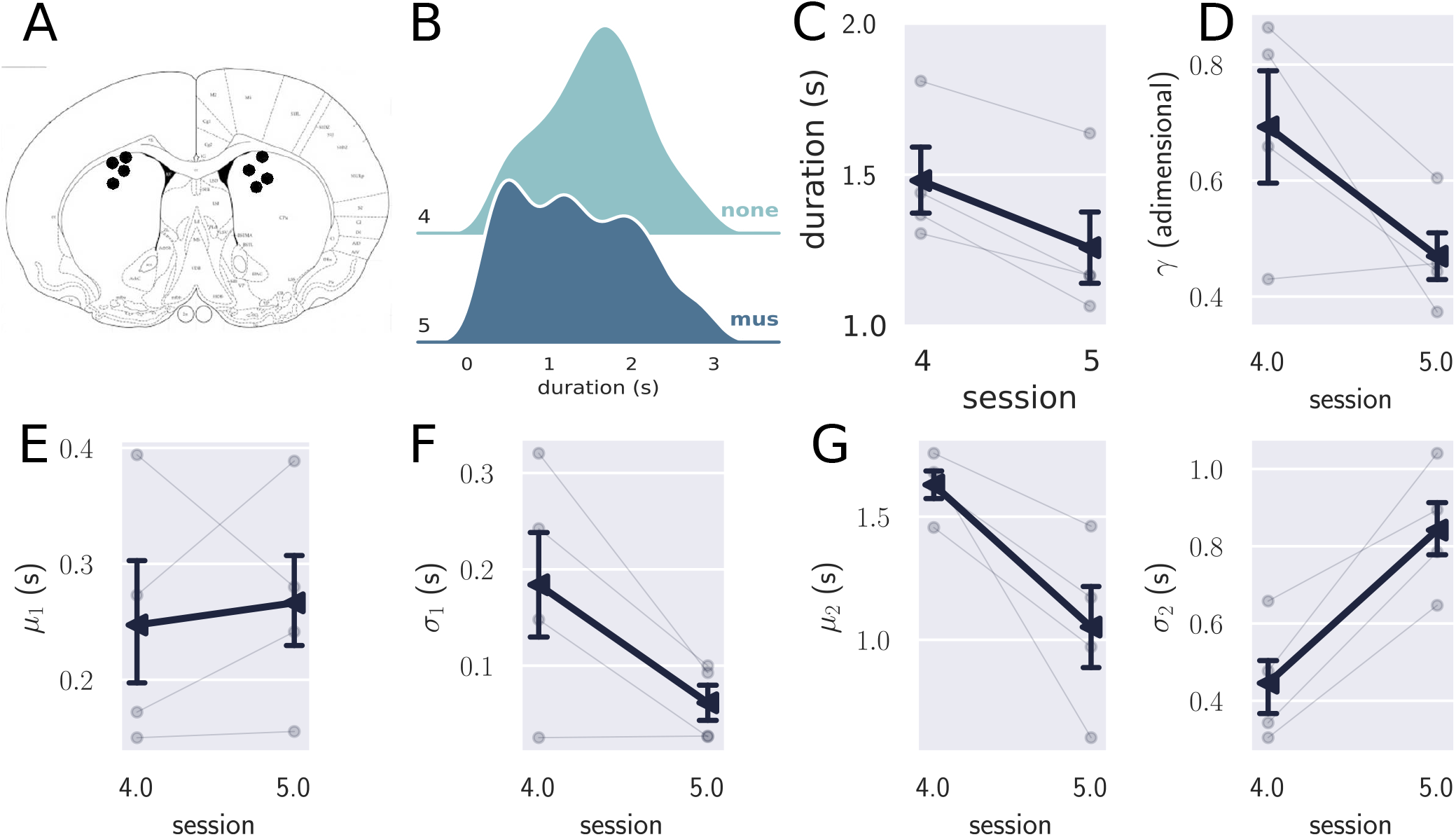
Inactivation of the STR in experienced animals. A) Histological sites for the pharmacological experiments in the STR. B) Comparison between the combined distribution of responses for experienced rats in session 4 (no drug) and session 5 (muscimol), showing that the temporally controlled responses were more variable when the STR was inactivated. C) Mean response duration for sessions four (no drug) and five (muscimol). D–H) Parameters obtained with the double Gaussian fit for sessions four and five. The variables *μ*_2_ and *σ*_2_ differed in these sessions, showing that the temporally controlled responses spread more with the inactivation of the STR.

In sum, the inactivation of STR led to a decrease in *μ*_2_ to a level comparable to that observed in the very first session of training, while *σ*_2_ increased to a level higher than that observed in session one. Such changes suggest a significant decrease in the time precision of the temporally-controlled responses. Interestingly, mean and standard variation of the premature responses (first Gaussian) did not change in the inactivation session, which suggests that the STR was more involved with the production of time-controlled than in the premature responses.

## Discussion

In this study, we investigated mPFC and STR role and activity while rats learned a temporal task. We tested the hypothesis that the time-related neuronal patterns classically found in trained animals should emerge especially in the mPFC as rats learned to correctly time their responses. Contrary to our hypothesis, results showed an initial involvement of the mPFC in the task (with a lower participation of the STR early on), but a diminishing role as the rats quickly became proficient in the task (with a higher participation of the STR after learning). These findings were consistent both from a decoding analysis of neural activity from mPFC and STR made early and late in the task, as well as from selective inactivation of those areas during learning.

The decoding analysis revealed a better performance early in the task for the mPFC compared to the STR neurons, but better decoding performance for the STR than mPFC cells later in the task. Correspondingly, pharmacological inactivation of mPFC showed that this structure was necessary for learning but not the expression of timed behavior, while the inactivation of STR after learning led to impairment of performance.

The decoder approach has been proposed to study the information encoding in different brain regions. Bakhurin and colleagues^19^ showed that these algorithms could quantify the amount of information encoded in different mice brain regions — the STR and the orbitofrontal cortex — in a Pavlovian conditioning task. They concluded that both regions encoded time, but the STR outperformed the orbitofrontal cortex. We cannot compare our results directly with those from Bakhurin et al. because they recorded from experienced (over-trained) animals (5 to 10 training sessions), they recorded from a different species, and they registered in a different area of the mPFC. However, on the second day of training, our results show that the STR outperforms the mPFC, a situation likely to remain stable since most learning happens on the first day of training^25^. Hence, it is possible that after the first day (or days) of training, the STR became even more reliable than the cortex in encoding the time of events, which agrees with results from Bakhurin and colleagues^19^.

Previous studies attempted to describe the collective reorganization of neuronal activity by recording in naive animals during their first day of learning^9, 17, 26^. Modi et. al^l9^ recorded from CA1 hippocampus neurons in a classical task — the trace eye-blink conditioning. They used calcium-imaging techniques to identify clusters of neurons sequentially firing during the acquisition. They reported a progressive increase in firing sequences within the trial, concomitantly to improvements in behavioral performance. They also used the a noise-correlation technique to infer common inputs to the network. Interestingly, the sequential firing observed in learners was preceded by a transient increase in noise correlations, which can be interpreted as a transitory increase in a common input to the CA1. Even though their behavioral task differs from ours, and the absence of mono-synaptic projections from mPFC to CA1^27^, these common transient effects seem an essential part of learning, and there might be a causal connection between them.

The decoding method proved to be useful in detecting changes in the network dynamics due to learning. Interestingly, the neuronal activity was quite organized from the very beginning, shown by a good decoding performance early on the first day of training. Such performance may relate to the fact that animals practiced to nose poke during the FR1 training phase. Hence, animals may have habituated with the action (nose poking), and the only new contingency introduced in the task was the wait time. In this sense, our results agree with previous results implicating the mPFC in top-down control of executive tasks, including behavioral switching learning^28–31^.

Our pharmacological results also provided causal evidence of the mPFC role in rats’ ability to time longer responses. They revealed a substantial impairment in learning when the mPFC was inactivated, but this effect did not last after the first session. During the second session, rats from group Muscimol significantly improved during training, revealing that after the first session, rats learned — at least partially — even with their mPFC inactivated. One possible explanation is that the pharmacological procedure only partially inactivated the mPFC, and the remaining active neurons provided some learning. Also, other cortical areas may underlie the initial learning phase of this task, hence producing the observed change in behavior. Finally, the number of converging projections to the STR^32^ makes it prone to detect environmental changes from multiple brain sources beyond the mPFC. Hence, the STR may provide a different action selection and, consequently, a different strategy to optimize reward rate^29, 33^.

Another result of the inactivation of the mPFC was that this region completely disengages from timing after learning. Inactivation of mPFC did not disturb performance if the rats had already learned the task. Narayanan et al.^34^ reported similar results in a different paradigm, a reaction time procedure. They showed that the inactivation of mPFC had an effect on the premature responses, but no effect on the time-controlled responses. Smith and collaborators also showed that the mPFC disengages from the task during habit formation^35^. However, these results are at odds with several results showing the critical role of the mPFC in timing^15, 23, 24, 36^. Particularly, results from different tasks show that this region’s inactivation impaired the timing precision but not accuracy^23, 24^. Then, since we dealt with a timing task, it seemed natural to assume the mPFC would be necessary for the expression of timed behavior. Such a hypothesis did not hold since inactivating the mPFC produced no detectable effect on experienced animals.

The mPFC’s disengagement interrogates why our results differ so antagonistically from those in the literature. One possible explanation has to do with the different time scale of our experiment. The 1.5 s interval used in our task is shorter than that used in most timing experiments. The interval of 1 s is usually referred to as the point of division between interval-timing and the millisecond range^37^. Our interval is slightly above one second, and yet our results regarding the mPFC role are quite distinct from experiments of more extended intervals^23, 24^. However, we cannot attribute the differences exclusively due to the interval duration. Xu et al.^15^ used similar intervals than ours — training animals to reproduce auditory stimuli either 1.5 or 2.5 s long — and found that neuronal activity scaled with the estimated interval. They also manipulated the temperature of the mPFC, which biased the responses in time, providing a causal link between the mPFC and the time estimation. We hypothesize that the difference between Xu and colleagues and our results rest on the fact that our task is self-timed, i.e. it does not rely or depend on external stimuli, which may give the task a more autonomic character, and reducing the number of required associative links to its execution.

The increasing involvement of the STR in the second day of training seen our electrophysiological experiments were expected since the known role of this region in the expression of timed responses^8,12,18^. However, our results give a timescale to this phenomenon, showing that the increase in time encoding happens already on the second day of training. Recently, Monteiro and collaborators^12^ showed a causal relationship between the dynamics within the STR and the decision based on timing in a temporal classification task, whose decision interval was 1.5 s. They manipulated the STR temperature showing an underestimation of time in increased temperatures and overestimation when the STR was cooled, consistent with the interpretation that the intrinsic dynamics of the STR underlies time estimation. When we inactivated the STR the timing performance was also disrupted, providing another causal link between the STR function and timing. Interestingly, the STR’s inactivation did not bias the response durations, only increased their variability. Such results suggest that other regions beyond the dorsomedial STR contribute to the timed responses. Regardless, our results point at the importance of the STR after the first day of training, deepening the view that the STR robustly underlies time estimation in a wide variety of timing modalities.

Finally, our results agree with recent neuronal recordings during learning of a new interval in a fixed-interval task28. They showed that the medial prefrontal cortex and striatum play different rules on temporal and behavior flexibility. While PFC was more likely related to the behavior flexibility, striatum was modulated by the temporal rule.

Overall, the our results provide a double dissociation between the role of the mPFC and the STR in a timing task. They also further our understanding of regions involved in time estimation and production, improving our knowledge of a taxonomy of time^38, 39^.

## Methods

### Subjects

The subjects were sixty one adult naive male Wistar rats 12 - 16 weeks old and weighing between 350 - 400g (purchased from the Federal University of São Paulo, Brazil). Eighteen animals were used in the STR pharmacological protocol, thirty five for mPFC pharmacological protocol, and eight at the electrophysiology protocol (four implanted in the mPFC and four in the mPFC and STR). Animals were housed individually in a 12 h light/dark cycle (lights on at 7 am). All procedures were conducted during the light cycle. Rats were gradually food-deprived to reach and maintain 85% of their free-feeding weight. Water was freely available. All experimental protocols were approved by our institutional animal care and use committee (CEUA-UFABC).

### Apparatus

Animals with chronic implants of electrode arrays were trained in an operant chamber developed in our laboratory, controlled by an Arduino Uno board. The box was built of acrylic plastic. On one wall, a nose poke was equipped with an infrared emitter-sensor beam, which interrupts when the rat inserts its snout. There was a bottle with a metallic nozzle on the left of the same wall, from which the rat can lick to obtain the reward. This nozzle connects to a touch-sensitive electronic device that counts the number of licks. A metallic plate controlled by the Arduino circuit moves up and down, blocking or releasing the rat access to the nozzle.

We used six MED-Associates operant chambers for the pharmacological procedure equipped with two levers, two cue lights (one above each lever), and a food cup on the front wall; more details are presented in Reyes et al.^25^. The rats executed an identical protocol as the rats trained in the electrophisiology boxes, except that the responses were produced in a lever. The response duration was measured from the time the rats pressed the lever to the moment they released it.

### Timing procedure

Animals implanted with electrodes were trained in our custom-made chambers to respond in the nose poke and receive access to three licks of 50% sucrose solution as a reward. Animals in the pharmacological experiments used the Med-Associates training chambers, responded on the left lever and received a 45 mg sugar pellet as a reward. Regardless of the chamber used, the behavioral procedure was the same. First, the animals were auto-shaped to nose poke (press a lever) in a fixed ratio 1 (FR1) schedule of reinforcement, in which the animal received access to the glucose solution (sugar pellet) after each response. Rats received 60 min daily sessions in this schedule until they responded 100 times in a single session. In the next session, animals started the timing procedure described in Reyes et al.^25^. Trials were self-initiated and self-ended by responding on the nose-poke (lever press). The animal had to nose poke (sustain the lever pressed) for at least 1.5 s to receive the reward: three licks in a 50% concentrated glucose solution (one 45 mg sugar pellet). The reward was only available after the animal self-ended the trial. Responses shorter than 1.5 s produced no consequences, and the rat could immediately start a new trial. If the animal did not consume the reward at the end of the trial, it could start a new trial. Rewards were available until the animal consumed them.

### Pharmacology

Bilateral cannulas were implanted in the striatum of eighteen rats. Three rats were removed from the experiments because they did not learn the FR1 after 5 session, and 11 were eliminated due to problems during drug injections or cannula placement. We also implanted bilateral cannulas on mPFC of thirty five rats (male Wistar). From this group one was removed because it did not learn the FR1 after 5 session, 15 were removed because of problems in drug injections or cannula placement. The stereotaxic coordinates were: mPFC AP +3.24 mm; ML = ±0.1 mm; DV = −3.3 mm; striatum: AP −2.3 mm; ML = −3 e +3 mm; DV = −1.5 mm. After one week of recovery from the surgery, the animals were food-restricted, and the experiments began. Infusion of muscimol or saline at 100 ng/0.5*μ*l were made 10 minutes before the beginning of the behavioral session. Injectors were inserted into the guide cannula, and 0.5 *μ*l of infusion fluid was delivered per site at 0.5*μ*l/min rate of infusion. After infusion was complete, the injector was held in place for 1 minute to allow fluid diffusion.

For the STR manipulation, we used a single injection after learning (fifth session), and for mPFC manipulation, we separated the animals into two groups. The group Muscimol received muscimol during the three first sessions, no infusion in the fourth session, and muscimol infusion again in the fifth session. In contrast, the group Saline received saline infusion in the three first sessions, no injection in session 4 and muscimol infusion in session five.

## Unit recordings and Data Analysis

The electrophysiological data for the mPFC (n=4) were obtained implanting each animal with a 32 electrodes array made by TDT (Tucker-Davis Technologies, EUA) in the right mPFC. Stereotaxic coordinates: AP = 2.52–4.68 mm; ML = 0.1–1.5 mm; DV = 3.3 mm. Simultaneous recordings from mPFC and striatum (n=4) were obtained with animals implanted with a 16 electrode arrays in the mPFC (AP = 2.52–4.68 mm; ML = 0.1–1.5 mm; DV = 3.3 mm) and a 16 electrode array in the striatum (AP = −0,23 mm; ML = −3,0 e +3,0 mm; DV = −1,5 mm). These electrodes were manufactured in our laboratory using omnetics connectors and 50 *μ*m tungsten microwires. The signal was recorded with a TDT system and digitized at 25 KHz in the first experiment. For the second experiment, the signal was recorded with an Open Ephys system. The raw signal was filtered between 1 to 5 kHz to select high-frequency events. A threshold of 2 standard deviations of the signal was used to select spikes. Finally, spike sorting was performed first online and then offline with the Open Sort TDT software. A PCA and waveform shape were used to spike sort, and then the clusters of spikes were discriminated by a semi-automatic system. First, we used a k-means method to select 3 clusters; then, the cluster was adjusted by visual inspection; noise clusters were excluded. Spike activity was analyzed for all cells that presented at a refractory period of at least 2 ms.

## Surgical and Histological procedures

Animals were anesthetized using a cocktail ketamine (100 mg/kg IF) and xylazine (10 mg/kg IP). The surgical level of anesthesia was maintained using ketamine supplementary doses of 0.1 ml when the surgical duration was above 40 minutes or it the animal presents responded to stimulus on the tail or feet. While the animal was on a superficial anesthetic plan, we started a sepsis protocol and accommodated the animal on the stereotaxic apparatus. When the animal was under a deep anesthetic plan, we made an incision. Craniotomies were drilled above the mPFC or striatum, from where the electrodes or cannulas were implanted. Also, four holes were drilled for skull screws, which were used to implant the ground wire e keep the implant’s stability. After the electrode placed into mPFC or STR, the craniotomy was closed with dental acrylic. Rats recovered for one week before the behavior training.

Following the experiment, rats were anesthetized and euthanized with an injection of 100 mg/kg of urethane. To map the electrode position, we used an electrical current to promote damage where the electrodes were implanted. Moreover, to map the cannula position, the animal received a microinjection of methylene blue dye. Then we transcardially perfused the rats with 4% formalin. Brains were post-fixed in a solution of 8% formalin; after 2 days, the brains were transferred to a 30% sucrose solution until they sunk to the bottom of the falcon tube. Next, they were frozen in isopentane at −80°C and sectioned in 25-micrometer slices in a cryostat. Brain slices were mounted on gelation-subbed slides and stained for cell bodies using Cresyl violet. Finally, the electrode and cannulae placements were defined by optical microscopy analysis.

## Data Analysis

### Electrophysiological Data

The neural activity analysis were performed in python scripts developed in our lab. To calculate the peri-event raster and histograms we computed the spiking activity from 0.5 s before the animals start (onset) a response (nose-poke or lever press) until the moment they they end the response (offset). Also, we calculated the peri-events for each neuron considering the neural activity aligned by the onset and by the offset of each trial. The peri-event histograms were calculated using bins of 100 ms, and were smoothed with a bandwidth of 100 ms.

### Multivariate Pattern Activity

We used separate multivariate analysis for the two groups of animals, one with recordings from mPFC and the other with simultaneous recordings from mPFC and STR. All analyses were performed on spiking data from all animals within groups, in a way that all registered cells were considered to be from an unique ”average” rat. We used the spikes from reinforced trials, i.e., trials longer than the reinforcement criterion (1.5 s). Trials were truncated at the criterion time and the average fire rate was calculated in bins of 100 ms. We set the number of trials from the ”average” rat as the smallest number of reinforced trials, such that all the cells were present in the trials used for classification. We used a multiclass linear discriminant analysis (LDA) as implemented in the sci-kit Python library, with least squares solver, automatically calculated shrinkage using Ledoit-Wolf lemma, no priors and the default number of components: min(number of classes - 1, number of features). Cross-validation was performed using 100 random folds. In each fold, training data consisted of activity from 10 bins (one bin for each class) from 80% of the trials. Bins from the same trial were always in either the train or test set. Training and testing data were normalized based on the median and inter-quartile range for each neuron, calculated over the training set. The output for each fold was the probability of each class, that was in turn averaged across folds to generate the confusion matrices. Performance measures were estimated in each fold using a Pearson correlation from the true class with the highest-probability class for each bin. At the group level, performance was compared to chance using a one-sample t-test. Performance as a function of learning was compared using unpaired t-test.

### Behavioral Data

All data were analyzed using Python routines developed in our laboratory. Statistical analysis was performed in Jasp^40^ and Pingouin^41^. Our dependent variable was response duration. For each animal, we constructed probability density diagrams of response duration. To test if the animals learned the task in the first session, we compared a group of trials from the beginning and the end of the session. For the electrophysiology and STR protocol, we used paired t-tests. For the PFC pharmacological protocol, we used a two-way ANOVA to compare groups and moments of the session.

As we showed in our previous work^25^, animals trained in the task frequently display a bimodal distribution, characterized by the persistence of very short responses interspersed with more timed responses — even after several sessions of training^42^. To better characterize bimodal distributions, we used a double Gaussian fit. We fitted the responses of each animal and condition separately, using as ordinary least squares method “curve fit” from the package “scipy.optimize”. The initial paramaters to start the fitting iteractions were: *γ* = 0.5, *μ*_1_ = 0.2, *σ*_1_ = 0.1, *μ*_2_ = 1, and *σ*_2_ = 0.5.

The double Gaussian probability density function was defined as *p*(*t*_*i*_) = *f*(*t*_*i*_)*/ξ*, where

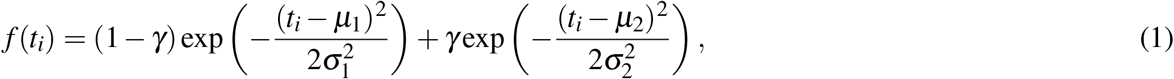

and the normalization term

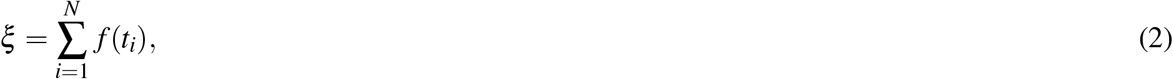

with time measured from 0 to 6 s in 0.1 s bins: *t*_*i*_ = *i* * 0.1 *s*, *i* = 0 ... 60.

At the group level, estimated parameters were compared using parametric tests, such as t-tests and ANOVA’s, indicated in each case.

## Acknowledgements

This work has bee partially supported by grants #2016/18914-7, #2016/05473-2, #2018/20277-0, São Paulo Research Foundation (FAPESP), and grant #430993/2016-1, National Council for Scientific and Technological Development (CNPq). Authors thank Nandakumar Narayanan for valuable feedback on the manuscript.

## Author contributions statement

MBR, GCT and EFO designed the experiment, and analyzed the data. GCT built the experimental setup for electrophysiology, and adapted the experimental procedure to the recordings. EFO set up the electro-phyisiological recordings. GCT recorded the activity in the mPFC and conducted all pharmacological experiments. EFO recorded the activity simultaneously in mPFC and STR. EUPV performed the machine learning analysis. MSC and AMC contributed to the experiment design, data analysis, and to the advance of the later versions of the manuscript. MBR and GCT wrote the first versions of the manuscript, revised by all authors.

## Additional information

The authors declare no competing interests.

